# Network Segregation in Aging Females and Evaluation of the Impact of Sex Steroid Hormones

**DOI:** 10.1101/2022.10.12.511918

**Authors:** Tracey H. Hicks, Thamires N. C. Magalhães, Hannah K. Ballard, T. Bryan Jackson, Sydney J. Cox, Jessica A. Bernard

**Affiliations:** Department of Psychological & Brain Sciences, Texas A&M University, College Station, TX, USA; Texas A&M Institute for Neuroscience, Texas A&M University, College Station, TX, USA

**Keywords:** network segregation, functional connectivity, sex hormones, aging

## Abstract

Males and females show differential patterns in connectivity in resting-state networks (RSNs) during normal aging, from early adulthood to late middle age. Age-related differences in network integration (effectiveness of specialized communication at the global network level) and segregation (functional specialization at the local level of specific brain regions) may also differ by sex. These differences may be due at least in part to endogenous hormonal fluctuation, such as that which occurs in females during midlife with the transition to menopause when levels of estrogens and progesterone drop markedly. A limited number of studies that have investigated sex differences in the action of steroid hormones in brain networks. Here we investigated how sex steroid hormones relate to age-network relationships in both males and females, with a focus on network segregation. Females displayed a significant quadratic relationship between age and network segregation for the cerebellar-basal ganglia and salience networks. In both cases, segregation was still increasing through adulthood, highest in midlife, and with a downturn thereafter. However, there were no significant relationships between sex steroid hormone levels and network segregation levels in females, and they did not exhibit significant associations between progesterone or 17β-estradiol and network segregation. Patterns of connectivity between the cerebellum and basal ganglia have been associated with cognitive performance and self-reported balance confidence in older adults. Together, these findings suggest that network segregation patterns with age in females vary by network, and that sex steroid hormones are not associated with this measure of connectivity in this cross-sectional analysis. Though this is a null effect, it remains critical for understanding the extent to which hormones relate to brain network architecture.

## 1. Introduction

Advanced age is associated with an overall decrease in the effectiveness of specialized communication at the global network level (ie, integration) and loss of functional specialization at the local level of specific brain regions (ie, segregation). That is, neuronal networks become less distinct with advanced age (1).

Age-related differences in the reorganization of functional connectivity and cognitive abilities may also differ by sex. In adulthood, sex differences in brain structure and function can be observed (2) and, similarly, sex differences in resting state networks (RSNs) have been reported. For example, studies of age differences have observed that males show increasing connectivity between-networks when compared to females (3–5). Males also exhibited more marked changes in default mode network (DMN) connectivity, especially in the posterior cingulate cortex (PCC), but showed smaller differences (and possibly increases) in connectivity to the lateral prefrontal regions of the fronto-parietal network (FPN) relative to females (4, 5). Females on the other hand showed smaller differences in DMN connectivity but showed greater decreases in FPN connectivity when compared to males.

Sex differences in network architecture have also been reported (6). Zhang and colleagues (2016) observed that female functional networks have significantly more connected nodes than males suggest an increase in network homogeneity in female brains. They also observed that the cerebellar nodes have a higher clustering coefficient and local efficiency for females (6). This adds evidence to their findings that the clustering coefficient and local efficiency in males are higher, while female connections are diffuse across lobes and the network is less modular. These results jointly support the notion that networks in female brains, compared to those in males, are more spatially distributed but with lower correlation strengths (6, 7). Overall, males and females showed differential patterns in connectivity in RSNs during normal aging, and from early adulthood to late middle age (4, 5, 7). However, these differences vary between studies with respect to regions and networks that are impacted.

Just as there are mixed results regarding sex differences with age in brain networks, there are also disagreements in the literature regarding differences in restingstate functional connectivity in the context of hormonal differences between biological sexes. To this point, there have been a limited number of studies that have investigated sex steroid hormones on brain networks. Further, there is currently disagreement among the rapidly expanding number of studies on the possible neuroprotective effects of sex hormones on cognitive and brain function more generally (8–10). Given that RSNs appear to be differentially impacted in males in females in later life, further exploration of the impact of sex steroid hormone levels on network architecture across adulthood stands to improve our understanding of underlying factors contributing to these differences. Specifically, it may be that sex hormone levels are related to the integration and segregation of neuronal networks (11).

In the brain, hormone receptors can be found across several regions. Post-mortem studies have found estrogen receptors in the hippocampus, claustrum, cerebral cortex, amygdala, hypothalamus, subthalamic nucleus, and thalamus (12–14). As for progesterone, a post-mortem study reported high concentrations of its receptors in the amygdala, hypothalamus, and cerebellum (14, 15). Testosterone exerts an early organizational effect on the development of the hypothalamus (16), the cerebral cortex (17), and the hippocampus (18) in addition to other brain structures. With respect to testosterone however, it is notable that it can interact not only with androgen receptors, but also with estradiol receptors (8). Therefore, it is also important to investigate the interactions that may occur between sex hormones and whether together they may be involved related to aging processes, as well as differences in neuronal networks (14). Testosterone, progesterone, and estrogens are present in both males and females, but their levels and production vary, mainly with respect to sex and age, though fluctuations in estrogens and progesterone occur across the female menstrual cycle as well (10).

The influence of sex hormones on functional networks is vital to our understanding of brain function and organization during periods of endogenous hormonal fluctuation, such as that which occurs in females during midlife with the transition to menopause when levels of estrogens and progesterone drop markedly (1, 9). Our study here aims to investigate how sex steroid hormones relate to age-network relationships in both males and females, with a focus on network segregation. As such, we predicted that there would be associations between network segregation and hormone levels, as well as interactions between hormones that may be affecting or enhancing the segregation of RSNs.

## 2. Methods

### 2.1. Study sample

One hundred and fifty-seven participants (total n=157) were enrolled as part of a larger study on aging. All participants underwent a battery of cognitive and motor tasks and during this assessment, the participants provided saliva samples for hormone quantification (for details about collection see *Hormone Quantification* below). After the behavioral visit the participants returned for a magnetic resonance imaging (MRI) session approximately two weeks later. However, due to unexpected delays related to the Covid- 19 pandemic, the time between the two sessions (39.0 days ± 21.4 days) varied between participants. For our analyses here, we focused only on the hormone and brain imaging data

Exclusion criteria were history of neurological disease, stroke, or formal diagnosis of psychiatric illness (e.g., depression or anxiety), contraindications for the brain imaging environment, and use of hormone therapy (HTh) or hormonal contraceptives (intrauterine device (IUD), possible use of continuous birth control (oral) and history of hysterectomy). These latter exclusions were made to evaluate impacts of normative endocrine aging on healthy adult females. For our analyses here we focused only on those with available neuroimaging data and hormonal assays. Thus, our final sample included 121 participants (55 males (age 57 ± 14.76) and 66 females (age 57 ± 12.17)). A flowchart showing the exclusions and determination of the final sample for analysis is presented below **(Figure 1)**.

**Figure 1.**
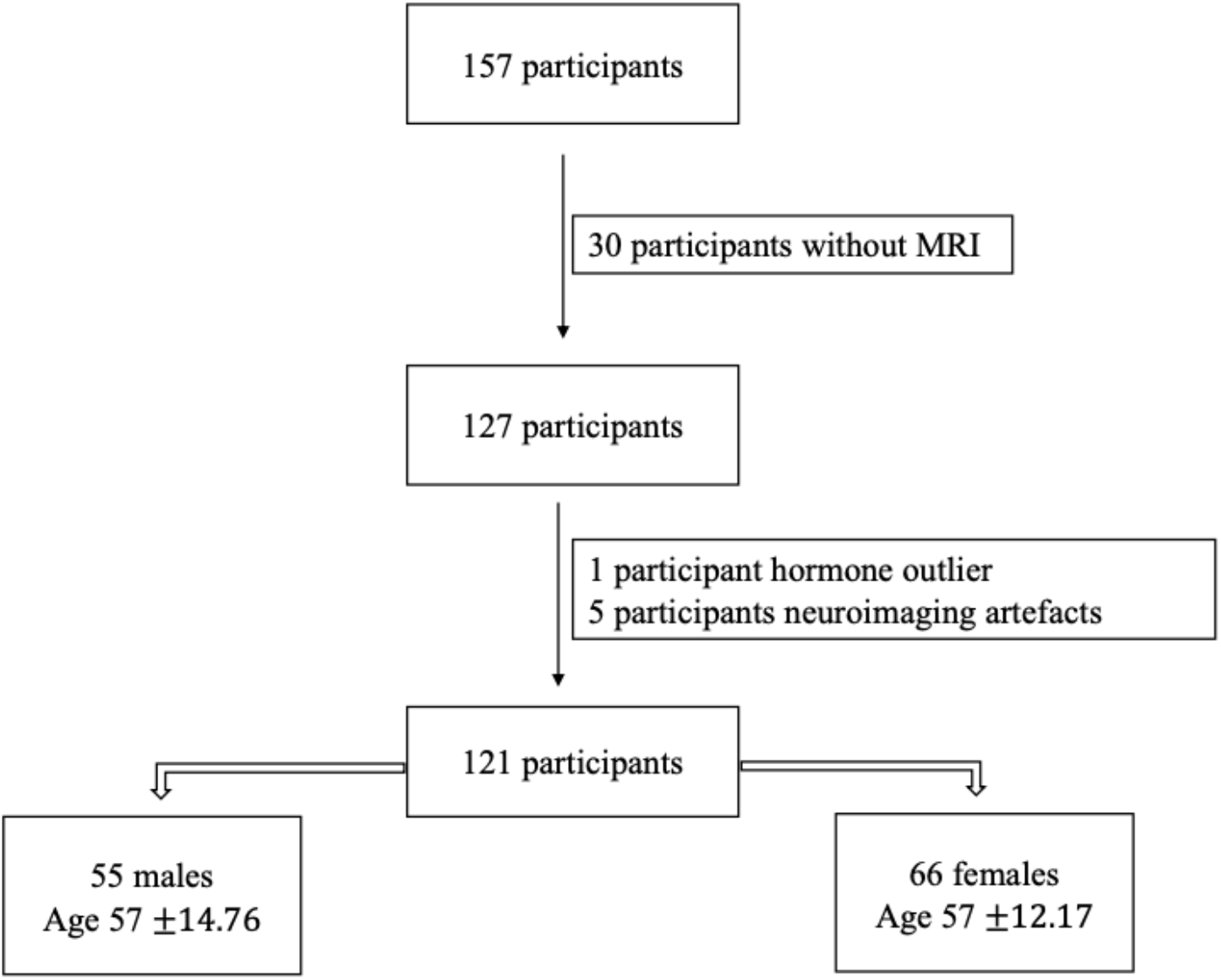
The initially collected number of participants and after following the exclusion criteria, the final number of our sample.

All study procedures were approved by the Institutional Review Board at Texas A&M University, and written informed consent was obtained from each participant prior to initiating any data collection.

### 2.2. Hormone quantification

For hormonal analyses, we followed the methodology used in recent work from our research group (31). For the sake of clarity and replicability, the methods reporting here matches what was reported in our recent work (31). Before collecting the saliva sample, participants were asked to refrain from consuming alcohol 24 hours prior and eating or drinking 3 hours prior to their first study session to avoid exogenous influences on hormone levels. Participants were also screened for oral disease or injury, use of substances such as nicotine or caffeine, and prescription medications that may impact the saliva pH and compromise samples. Participants were asked to rinse their mouth with water 10 minutes prior to providing a saliva sample to clear out any residue.

Samples were then collected in pre-labeled cryovials provided by Salimetrics (https://salimetrics.com/saliva-collection-training-videos/) using the passive drool technique. For our study, participants were asked to supply 1mL of saliva, after which samples were immediately stored in a −80° Celsius bio-freezer for stabilization. Assays were completed by Salimetrics to quantify 17*β*-estradiol, progesterone, and testosterone levels for each participant. The amount of saliva collected was sufficient to detect 17β- estradiol at a high sensitivity threshold of 0.1 pg/mL (19), along with 5.0 pg/mL and 1.0 pg/mL thresholds for progesterone and testosterone, respectively.

The protocol used by Salimetrics includes two repetitions of each assay; thus, the values used in our analyses represent an average of both repetitions. A few samples were insufficient in quantity and were unable to be properly assayed (n = 3; 2 progesterone, 1 testosterone). The intra-assay coefficient of variability for our hormone samples was 0.15 for 17*β*-estradiol, 0.11 for progesterone, and 0.07 for testosterone. This non-invasive method is adequate for precisely measuring reproductive hormones. Salivary measurements are strongly correlated with blood-derived measurements to index sex hormone levels (17β-estradiol: r = 0.80; progesterone: r = 0.80; testosterone: r = 0.96, recovered from https://salimetrics.com/analyte/salivary-estradiol/).

### 2.3. Imaging acquisition

Participants underwent structural and resting-state MRI using a Siemens Magnetom Verio 3.0 Tesla scanner and a 32-channel head coil. For structural MRI, we collected a high-resolution T1-weighted 3D magnetization prepared rapid gradient multi-echo (MPRAGE) scan (repetition time (TR) = 2400 ms; acquisition time = 7 minutes; voxel size = 0.8 mm^3^) and a high-resolution T2-weighted scan (TR = 3200 ms; acquisition time = 5.5 minutes; voxel size = 0.8 mm^3^), each with a multiband acceleration factor of 2. For resting-state imaging, we administered four blood-oxygen level dependent (BOLD) functional connectivity (fcMRI) scans with the following parameters: multiband factor of 8, 488 volumes, TR of 720 ms, and 2.5 mm^3^ voxels. Each fcMRI scan was 6 minutes in length for a total of 24 minutes of resting-state imaging, and scans were acquired with alternating phase encoding directions (i.e., two anterior to posterior scans and two posteriors to anterior scans). During the fcMRI scans, participants were asked to lie still with their eyes open while fixating on a central cross. In total, the acquisition of images takes about 45 minutes, including a 1.5-minute localizer.

Scanning protocols were adapted from the multiband sequences developed by the Human Connectome Project (HCP) (20) and the Center for Magnetic Resonance Research at the University of Minnesota to facilitate future data sharing and reproducibility.

#### 2.3.1. Imaging processing

##### 2.3.1.1. Pre-processing

Images were converted from DICOM to NIFTI and organized into a Brain Imaging Data Structure (BIDS, version 1.6.0) format via the latest docker container version of bidskit (version 2021.6.14, https://github.com/jmtyszka/bidskit). Using the split tool distributed with the FMRIB Software Library (FSL) package (21), a single volume was extracted from two oppositely encoded BOLD images to estimate B_0_ field maps. Next, fMRIPrep (version 20.2.3, https://fmriprep.org/) was used to preprocess anatomical and functional images. The fMRIPrep preprocessing pipeline includes basic steps such as co-registration, normalization, unwarping, noise component extraction, segmentation, and skull-stripping.

While basic pre-processing was performed in the fMRI preparation, we also completed remaining steps in the Conn toolbox, version 21a (22). We used the default preprocessing pipeline, which consists of realignment and unwarping with motion correction, centering to (0, 0, 0) coordinates, slice-timing correction, outlier detection using a 95^th^ percentile threshold and the Artifact Rejection Toolbox (ART), segmentation of grey matter, white matter, and cerebrospinal fluid, normalization to MNI space, and spatial smoothing with a 5 mm full width at half-maximum (FWHM) Gaussian kernel. A band-pass filter of 0.008-0.099 Hz was applied to denoise data. The threshold for global-signal z-values was set at 3, while the motion correction threshold was set at 0.5 mm. After being de-spiked during denoising to adhere to the global mean, 6-axis motion data and frame-wise outliers were included as first-level covariates.

##### 2.3.1.2. ROI selection

In our study, we used the same ROIs selection used by Ballard and colleagues (2022), in work previously completed by our group (23). The MNI coordinates for each cortical node retrieved from Cassady et al. (2019) (24) (originally derived from Power et al. (2011) (25)) were selected by our group. Further, we included 20 subcortical nodes of the (extracted from Hausman et al. (2020), (26)) for the cerebellar-basal ganglia network (27–29). Cerebellar seeds were determined via the SUIT atlas (28, 29). Our final set of ROIs contained 234 nodes across 11 networks (10 cortical, 1 subcortical, see **Table 1** for a list of included networks). MNI coordinates for each node were translated to voxel coordinates, which were subsequently used to create spherical seeds with 3.5 mm diameters in FSL (21). These seeds were then treated as ROIs.

**Table 1.** Network Abbreviation Key.

| Abbreviation | Network |
| --- | --- |
| Au | Auditory |
| CBBG | Cerebellar-basal ganglia |
| COTC | Cingulo-Opercular Task Control |
| DA | Dorsal Attention |
| DM | Default Mode |
| FPTC | Fronto-Parietal Task Control |
| Sa | Salience |
| SSH | Sensory Somatomotor Hand |
| SSM | Sensory Somatomotor Mouth |
| Vi | Visual |
| VA | Ventral Attention |

First-level ROI-to-ROI relationships were evaluated with a bivariate correlation approach; those correlations are needed to calculate the variables within and between the network for each subject which are then used in the network segregation equation. Correlation values were transformed into z-values via Fisher’s r-to-z conversion (30). Corrections for multiple comparisons were applied during statistical analyses.

##### 2.3.1.3. Network segregation equation

For the analysis of network segregation, we again followed our previous work reported by Ballard and colleagues (2022) (23, 31) and based off analyses initially conducted by Chan et al. (2014) (32). Network segregation values were determined using **Equation 1** below. In the formula, *<u>z</u>_w_* corresponds to the mean correlation between ROIs within an individual network, and *<u>z</u>_b_* represents the mean correlation between ROIs of an individual network and all remaining ROIs of other networks. Group-level analyses were performed with a voxel threshold of p < .001 and cluster threshold, FDR-corrected, of p < .05.

**Equation 1.** Network segregation values were determined using this formula.

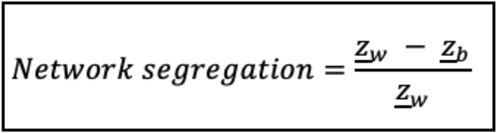

### 2.4. Statistical analysis

We first sought to ascertain sex differences in hormone levels within our sample.

Analyses of variance (ANOVAs) were conducted to determine sex differences in hormone levels (i.e., estradiol, progesterone, and testosterone separately). ANOVAs were completed using the anova function from the default ‘stats’ package in R (v4.0.5, R Core Team, 2021) which determined beta coefficients, degrees of freedom, F-values, and p-values; the sjstats and pwr packages were used to compute **η**^2^ and partial **η**^2^ values (v0.18.1, (33)).

As our primary area of interest lies within the impact of fluctuating hormones on female brain network segregation, our main analyses investigated females only. However, exploratory analyses evaluated males and all participants combined which are included in the supplement.

To explore the unique associations between hormone levels (i.e., estradiol, progesterone, and testosterone separately) and each network of interest in females, linear regressions were performed in which hormone levels served as the predictor and network segregation as the outcome. These linear regressions were also conducted in exploratory analyses with males and all participants collapsing the two sexes (Supplementary Tables I-VI). The lm function from the default ‘stats’ package in R (v4.0.5, R Core Team, 202; (34)) determined beta coefficients, degrees of freedom, F-values, p-values, R^2^, and adjusted R^2^. False discovery rate (FDR) correction was applied to account for multiple comparisons (i.e., number of networks examined) using the FSA package in R exclusively on results with a *p-value* ≤.05 (FSA v0.9.3, (35)). Linear regressions with hormone level interactions (i.e., estradiol*progesterone, estradiol*testosterone, and progesterone*testosterone) as the predictor variables explored the relationship between combined hormone levels and network segregation in females. Exploratory analyses investigated the same hormone level interactions in males and all participants together (***Supplement material*** **Tables 12-13**). FDR correction was applied as previously described (FSA v0.9.3, (35)).

Associations between network segregation and age were evaluated via linear regression with age as the predictor and network connectivity as the outcome in females; males and all participants together were run as exploratory analyses (***Supplementary material*** **Tables 12-13**). We conducted similar regressions with quadratic age (Age + I(Age^2)) as the predictor to investigate whether network segregation demonstrated better fit with a quadratic function rather than linear across middle-to advanced-age adults, given prior work suggesting non-linear relationships between brain system segregation (32) as well as brain volume (36) and age. Quadratic regressions for males and all participants were run as exploratory analyses (***Supplementary material*** **Tables 9-10**). Linear and quadratic regressions were performed using the lm function from the default ‘stats’ package in R (v4.0.5, R Core Team, 2021) which determined beta coefficients, degrees of freedom, F-values, p-values, R^2^, and adjusted R^2^. FDR correction was applied as previously described to account for multiple comparisons (i.e., number of networks examined) (FSA v0.9.3, (35)). Akaike’s An Information Criterion (AIC) (37) compared fit between linear and quadratic models with a requirement that model value must differ by 10 to be considered a superior fit (38, 39). AIC was calculated by the default ‘stats’ package in R (v4.0.5, R Core Team, 2021).

The stargazer package (v5.2.2; Hlavac, 2018) was used to create Tables II–VII. The ggplot2 package was used to create linear and quadratic plots (Figures 1-5; v3.3.3; (40)). All figures were created using a colorblind friendly palette via RColorBrewer (41).

**Table 2.** This table presents results from linear regressions for estradiol and network segregation in females. Raw *p*-values are listed, and FDR correction was only performed if raw *p* -value was <.05.

| Female Estradiol Linear Associations with Network Segregation |  |  |  |  |  |  |
| --- | --- | --- | --- | --- | --- | --- |
| | Estradiol $\beta$<br>coefficient | Raw $p$-value | R<sup>2</sup> | Adjusted R<sup>2</sup> | Residual Std.<br>Error (df = 55) | F Statistic<br>(df = 1; 55) |
| **Au** | -0.008 | 0.738 | 0.002 | -0.016 | 0.103 | 0.114 |
| **CBBG** | 0.039 | 0.313 | 0.019 | 0.001 | 0.157 | 1.037 |
| **COTC** | 0.914 | 0.385 | 0.014 | -0.004 | 4.27 | 0.769 |
| **DA** | -0.023 | 0.392 | 0.013 | -0.005 | 0.109 | 0.746 |
| **DM** | 0.018 | 0.625 | 0.004 | -0.014 | 0.146 | 0.242 |
| **FPTC** | 0.015 | 0.646 | 0.004 | -0.014 | 0.129 | 0.214 |
| **Sa** | 0.038 | 0.259 | 0.023 | 0.005 | 0.138 | 1.306 |
| **SSH** | -0.008 | 0.761 | 0.002 | -0.016 | 0.111 | 0.094 |
| **SSM** | -0.024 | 0.205 | 0.029 | 0.011 | 0.077 | 1.646 |
| **Vi** | -0.013 | 0.705 | 0.003 | -0.015 | 0.137 | 0.146 |
| **VA** | -0.022 | 0.731 | 0.002 | -0.016 | 0.26 | 0.120 |

**Table 3.** This table presents results from linear regressions for progesterone and network segregation in females. Raw *p*-values and FDR corrected *p*-values are included. There were no significant findings after FDR correction.

| Female Progesterone Linear Associations with Network Segregation |  |  |  |  |  |  |  |
| --- | --- | --- | --- | --- | --- | --- | --- |
| | Progesterone<br>$\beta$ coefficient | Raw *p*-value | FDR<br>Corrected *p*-<br>value | R<sup>2</sup> | Adjusted<br>R<sup>2</sup> | Residual<br>Std. Error<br>(df = 59) | F Statistic<br>(df = 1; 59) |
| **Au** | 0.0001 | 0.575 | 0.926 | 0.005 | -0.011 | 0.107 | 0.318 |
| **CBBG** | 0.0003 | 0.176 | 0.484 | 0.031 | 0.014 | 0.166 | 1.880 |
| **COTC** | 0.015 | 0.010 | 0.110 | 0.109 | 0.094 | 3.873 | 7.247 |
| **DA** | 0.0001 | 0.674 | 0.926 | 0.003 | -0.014 | 0.107 | 0.179 |
| **DM** | 0.0002 | 0.255 | 0.561 | 0.022 | 0.005 | 0.143 | 1.323 |
| **FPTC** | 0.0004 | 0.049 | 0.270 | 0.064 | 0.049 | 0.123 | 4.067 |
| **Sa** | 0.0003 | 0.102 | 0.374 | 0.045 | 0.029 | 0.134 | 2.764 |
| **SSH** | 0.0001 | 0.711 | 0.926 | 0.002 | -0.015 | 0.116 | 0.139 |
| **SSM** | 0.00001 | 0.926 | 0.926 | 0.0002 | -0.017 | 0.082 | 0.009 |
| **Vi** | 0.00004 | 0.850 | 0.926 | 0.001 | -0.016 | 0.133 | 0.036 |
| **VA** | -0.0001 | 0.881 | 0.926 | 0.0004 | -0.017 | 0.246 | 0.023 |

**Table 4.**
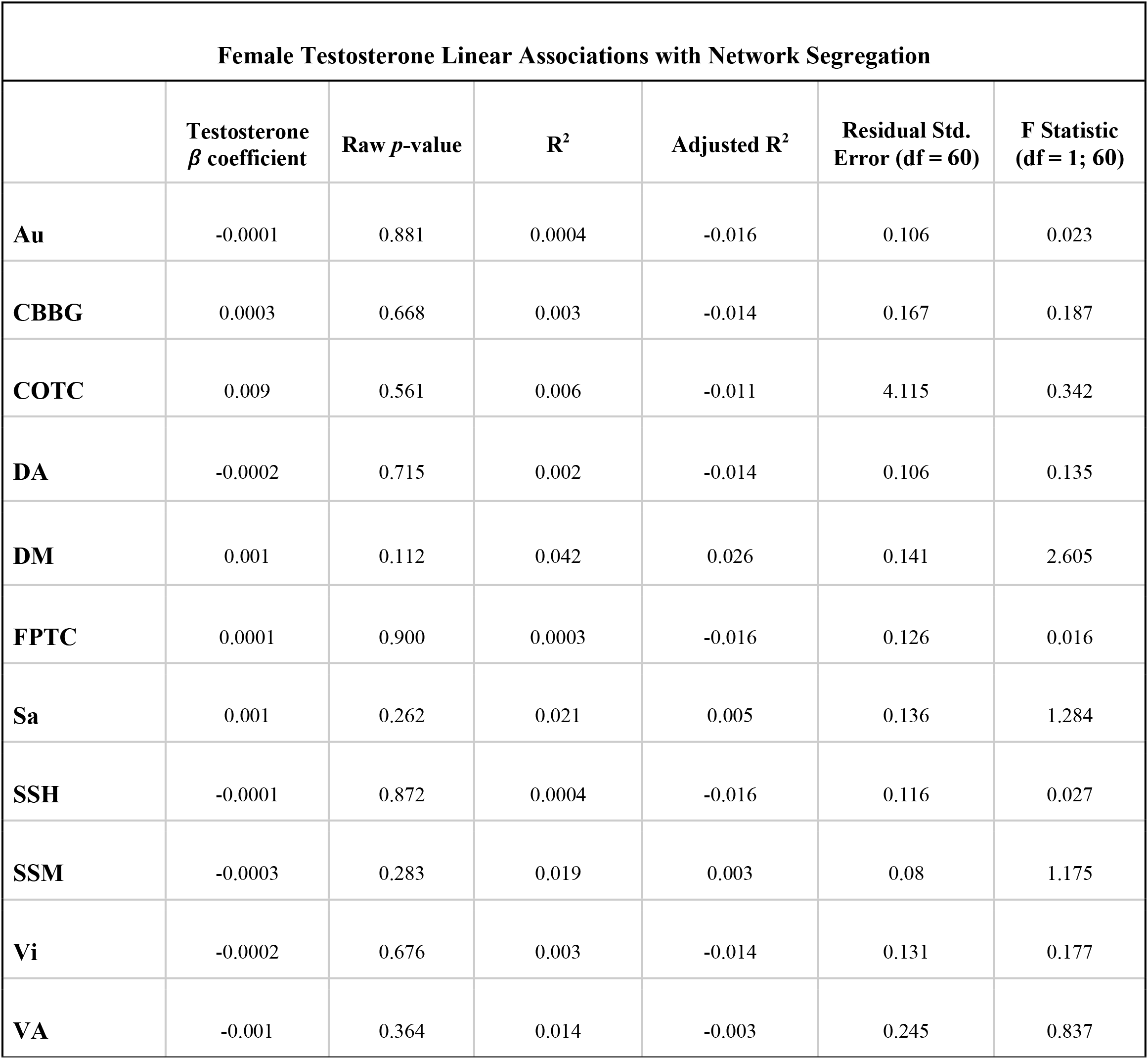
This table presents results from linear regressions for testosterone and network segregation in females. Raw *p*-values are listed, and FDR correction was only performed if raw *p* -value was <.05.

| Female Testosterone Linear Associations with Network Segregation |  |  |  |  |  |  |
| --- | --- | --- | --- | --- | --- | --- |
| | Testosterone $\beta$ coefficient | Raw $p$-value | $R^2$ | Adjusted $R^2$ | Residual Std. Error (df = 60) | F Statistic (df = 1; 60) |
| **Au** | -0.0001 | 0.881 | 0.0004 | -0.016 | 0.106 | 0.023 |
| **CBBG** | 0.0003 | 0.668 | 0.003 | -0.014 | 0.167 | 0.187 |
| **COTC** | 0.009 | 0.561 | 0.006 | -0.011 | 4.115 | 0.342 |
| **DA** | -0.0002 | 0.715 | 0.002 | -0.014 | 0.106 | 0.135 |
| **DM** | 0.001 | 0.112 | 0.042 | 0.026 | 0.141 | 2.605 |
| **FPTC** | 0.0001 | 0.900 | 0.0003 | -0.016 | 0.126 | 0.016 |
| **Sa** | 0.001 | 0.262 | 0.021 | 0.005 | 0.136 | 1.284 |
| **SSH** | -0.0001 | 0.872 | 0.0004 | -0.016 | 0.116 | 0.027 |
| **SSM** | -0.0003 | 0.283 | 0.019 | 0.003 | 0.08 | 1.175 |
| **Vi** | -0.0002 | 0.676 | 0.003 | -0.014 | 0.131 | 0.177 |
| **VA** | -0.001 | 0.364 | 0.014 | -0.003 | 0.245 | 0.837 |

**Table 5.** This table exhibits results from linear regressions for hormone interactions and network segregation in females. Raw *p*-values and FDR corrected *p*-values are included. FDR correction was only performed if raw *p* -value was <.05. There were no significant findings after FDR correction.

| Female Hormone Level Interactions with Network Segregation |  |  |  |  |  |  |  |  |  |  |  |
| --- | --- | --- | --- | --- | --- | --- | --- | --- | --- | --- | --- |
| | Estradiol by Progesterone (E*P) $\beta$ coefficient | Raw E*P $p$-value | Estradiol by Testosterone (E*T) $\beta$ coefficient | Raw E*T $p$-value | Progesterone by Testosterone (P*T) $\beta$ coefficient | Raw P*T $p$-value | FDR Corrected $p$-value for Estradiol by Testosterone (E*T) | R<sup>2</sup> | Adjusted R<sup>2</sup> | Residual Std. Error (df = 48) | F Statistic (df = 6; 48) |
| **Au** | 0.0003 | 0.669 | -0.003 | 0.072 | 0.00001 | 0.360 | 0.264 | 0.096 | -0.017 | 0.103 | 0.851 |
| **CBBG** | 0.001 | 0.131 | -0.003 | 0.167 | 0.00001 | 0.760 | 0.359 | 0.133 | 0.024 | 0.155 | 1.225 |
| **COTC** | -0.029 | 0.245 | -0.056 | 0.315 | 0.001 | 0.111 | 0.429 | 0.182 | 0.079 | 4.087 | 1.776 |
| **DA** | -0.0005 | 0.489 | -0.001 | 0.511 | 0.00001 | 0.507 | 0.511 | 0.05 | -0.069 | 0.111 | 0.422 |
| **DM** | 0.0002 | 0.792 | -0.002 | 0.333 | 0 | 0.856 | 0.429 | 0.085 | -0.029 | 0.148 | 0.747 |
| **FPTC** | -0.0002 | 0.786 | -0.002 | 0.196 | 0.00001 | 0.496 | 0.359 | 0.135 | 0.027 | 0.126 | 1.248 |
| **Sa** | 0.0001 | 0.940 | -0.002 | 0.359 | 0.00002 | 0.363 | 0.429 | 0.072 | -0.044 | 0.14 | 0.619 |
| **SSH** | 0.0002 | 0.755 | -0.003 | 0.035 | 0.00003 | 0.088 | 0.193 | 0.133 | 0.025 | 0.111 | 1.23 |
| **SSM** | 0.0001 | 0.766 | -0.002 | 0.028 | 0.00002 | 0.137 | 0.193 | 0.157 | 0.051 | 0.076 | 1.485 |
| **Vi** | -0.001 | 0.125 | 0.002 | 0.390 | 0.00001 | 0.711 | 0.429 | 0.059 | -0.059 | 0.142 | 0.497 |
| VA | 0.002 | 0.307 | -0.005 | 0.127 | 0.0000<br>1 | 0.691 | 0.349 | 0.072 | -0.044 | 0.26 | 0.617 |

**Table 6.** This table exhibits results from linear regressions for age and network segregation in females. Raw *p*-values and FDR corrected *p*-values are included. There were no significant findings after FDR correction.

| Female Linear Age Associations with Network Segregation |  |  |  |  |  |  |  |
| --- | --- | --- | --- | --- | --- | --- | --- |
| | Age $\beta$<br>coefficient | Raw $p$-value | FDR<br>Corrected $p$-<br>value | $R^2$ | Adjusted<br>$R^2$ | Residual<br>Std. Error<br>(df = 64) | F Statistic<br>(df = 1; 64) |
| **Au** | -0.002 | 0.024 | 0.088 | 0.078 | 0.064 | 0.105 | 5.414 |
| **CBBG** | -0.005 | 0.01 | 0.077 | 0.099 | 0.085 | 0.168 | 7.061 |
| **COTC** | -0.082 | 0.044 | 0.103 | 0.062 | 0.047 | 3.918 | 4.231 |
| **DA** | -0.001 | 0.311 | 0.342 | 0.016 | 0.001 | 0.106 | 1.043 |
| **DM** | -0.002 | 0.18 | 0.283 | 0.028 | 0.013 | 0.146 | 1.843 |
| **FPTC** | -0.003 | 0.047 | 0.103 | 0.06 | 0.046 | 0.126 | 4.114 |
| **Sa** | -0.004 | 0.014 | 0.077 | 0.092 | 0.078 | 0.141 | 6.466 |
| **SSH** | -0.001 | 0.276 | 0.337 | 0.019 | 0.003 | 0.113 | 1.210 |
| **SSM** | -0.001 | 0.065 | 0.119 | 0.052 | 0.037 | 0.077 | 3.529 |
| **Vi** | -0.001 | 0.357 | 0.357 | 0.013 | -0.002 | 0.13 | 0.862 |
| **VA** | -0.003 | 0.252 | 0.337 | 0.021 | 0.005 | 0.241 | 1.341 |

**Table 7.** This table exhibits results from quadratic regressions for age and network segregation in females. Raw *p*-values and FDR corrected *p*-values are included. Asterisks indicate significance at *p*<.05* for FDR corrected values. Only FDR corrected values are interpreted as significant.

| Female Quadratic Age Associations with Network Segregation |  |  |  |  |  |  |  |
| --- | --- | --- | --- | --- | --- | --- | --- |
| | Quadratic $\beta$<br>coefficient | Raw $p$-value | FDR<br>Corrected $p$-<br>value | $R^2$ | Adjusted<br>$R^2$ | Residual<br>Std. Error<br>(df = 63) | F Statistic<br>(df = 2; 63) |
| **Au** | -0.0002 | 0.014 | 0.051 | 0.164 | 0.138 | 0.1 | 6.202 |
| **CBBG** | -0.0003 | 0.002 | 0.011* | 0.241 | 0.217 | 0.155 | 10.020 |
| **COTC** | -0.002 | 0.391 | 0.430 | 0.073 | 0.044 | 3.925 | 2.482 |
| **DA** | -0.0001 | 0.071 | 0.112 | 0.066 | 0.036 | 0.104 | 2.23 |
| **DM** | -0.0002 | 0.043 | 0.095 | 0.09 | 0.061 | 0.142 | 3.109 |
| **FPTC** | -0.0002 | 0.058 | 0.106 | 0.113 | 0.085 | 0.124 | 4.011 |
| **Sa** | -0.0003 | 0.0005 | 0.011* | 0.255 | 0.232 | 0.129 | 10.806 |
| **SSH** | -0.0002 | 0.035 | 0.095 | 0.086 | 0.057 | 0.11 | 2.977 |
| **SSM** | -0.0001 | 0.16 | 0.210 | 0.082 | 0.053 | 0.077 | 2.808 |
| **Vi** | -0.0001 | 0.172 | 0.210 | 0.042 | 0.012 | 0.129 | 1.393 |
| **VA** | -0.0001 | 0.578 | 0.578 | 0.025 | -0.006 | 0.242 | 0.820 |

## 3. Results

### 3.1. Hormone Levels by Sex

An ANOVA revealed significant sex differences in testosterone when accounting for age (F (1,111) = 79.496, *p<.001*; Figure 1), that is testosterone levels were significantly lower in females. ANOVAs did not reveal significant sex differences in estradiol or progesterone levels (F (1,105) = 3.318, *p* = .*071*, η^2^= .031), and (F (1,109) = 3.270, *p*= .*073*, η^2^= .029), respectively; **Figure 2**).

**Figure 2.**
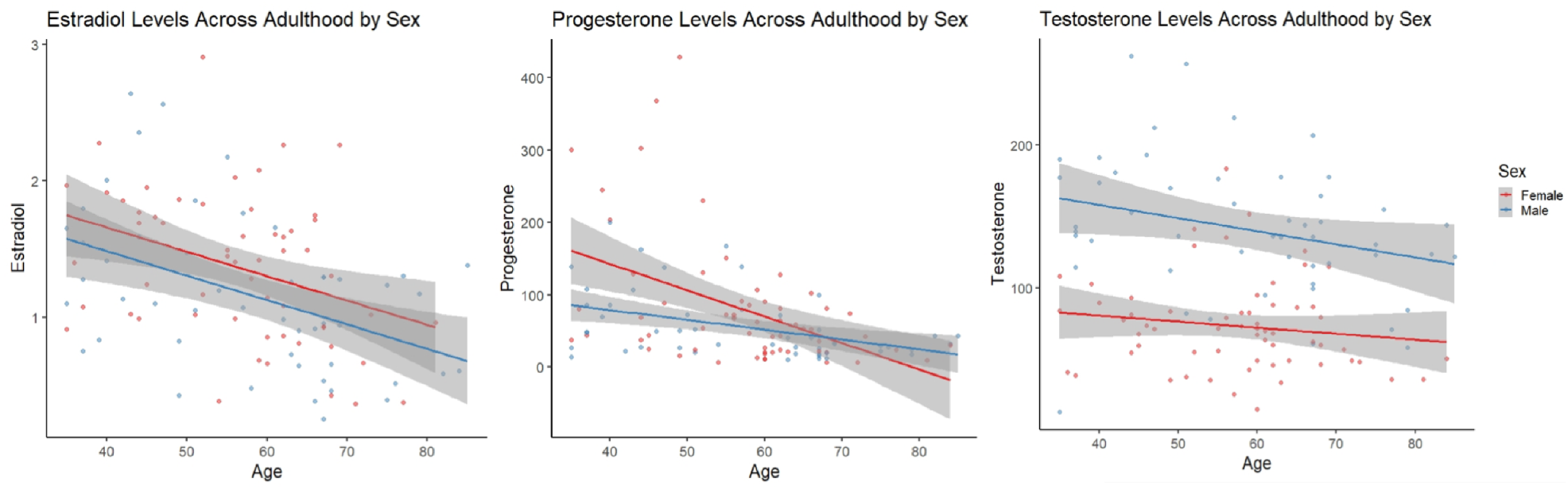
Linear scatter plots demonstrate estradiol levels in females and males across adulthood. The gray superimposed on each colored line depicts the 95% confidence interval in each sex. Estradiol levels (left) were not significantly different by sex when accounting for age (*p* = .071). Progesterone levels (middle) were not significantly different by sex when accounting for age (*p*= .073). Testosterone levels (right) were significantly different by sex when accounting for age (*p*< .001).

### 3.2. Network Segregation and Hormone levels in Females

When examining females alone, network segregation was not significantly associated with progesterone, estradiol, or testosterone in females after FDR correction (see **Tables 2–4**). Similar exploratory analyses were conducted with hormone levels (i.e., estradiol, progesterone, and testosterone; respectively) across participants and in males and all participants (see Supplementary **Tables 1–3**).

### 3.3. Interactions Between Hormone Levels and Network Segregation in Females

Regressions evaluating combined effects (interactions) of hormone levels (i.e., Estradiol*Progesterone, Estradiol*Testosterone, and Progesterone*Testosterone) as the predictor and network segregation as the outcome were not significant (see **Table 5**). Similar analyses were conducted in males and all participants (see Supplementary **Tables 12-13**). Notably, though the 17*β*-estradiol*testosterone interaction was associated with segregation in both the sensory somatomotor hand and mouth networks, these did not survive FDR correction.

### 3.4. Linear and Quadratic Associations Between Age and Network Segregation

Network segregation was evaluated in females across the adult lifespan with respect to age. Linear regressions with age as the predictor and network segregation as the outcome revealed no significant associations after corrections for multiple comparisons (see **Table 6**).

However, in females, regressions with quadratic age as the predictor and network segregation as the outcome, demonstrated significant associations in the cerebellar-basal ganglia (CBBG) (F (2, 63) = 10.020, raw *p* = .002, FDR adjusted *p* = .011) and salience (Sa) (F (2, 63) = 10.806, raw *p* < .001, FDR adjusted *p* = .011) networks (**Figures 3 and 4**; **Table 7**). There were no additional significant associations in network segregation and quadratic age for females or males as revealed in our exploratory analyses (detailed results are presented in **Table 7**).

**Figure 3.**
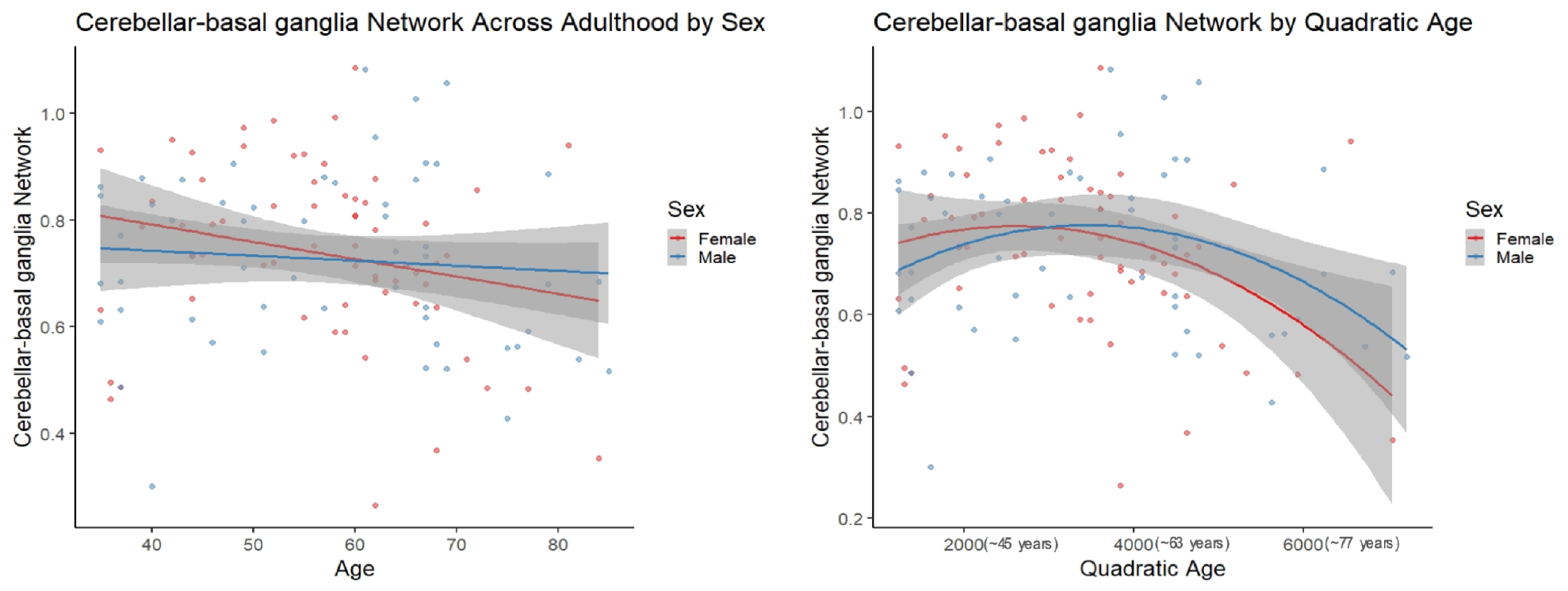
Scatter plots demonstrate the linear (left) and quadratic (right) relationship between Cerebellar-basal ganglia network connectivity and age. Parentheses next to quadratic values (right) indicate the approximate square root of the quadratic value for interpretative purposes. The gray superimposed on each colored line depicts the 95% confidence interval in each sex.

**Figure 4.**
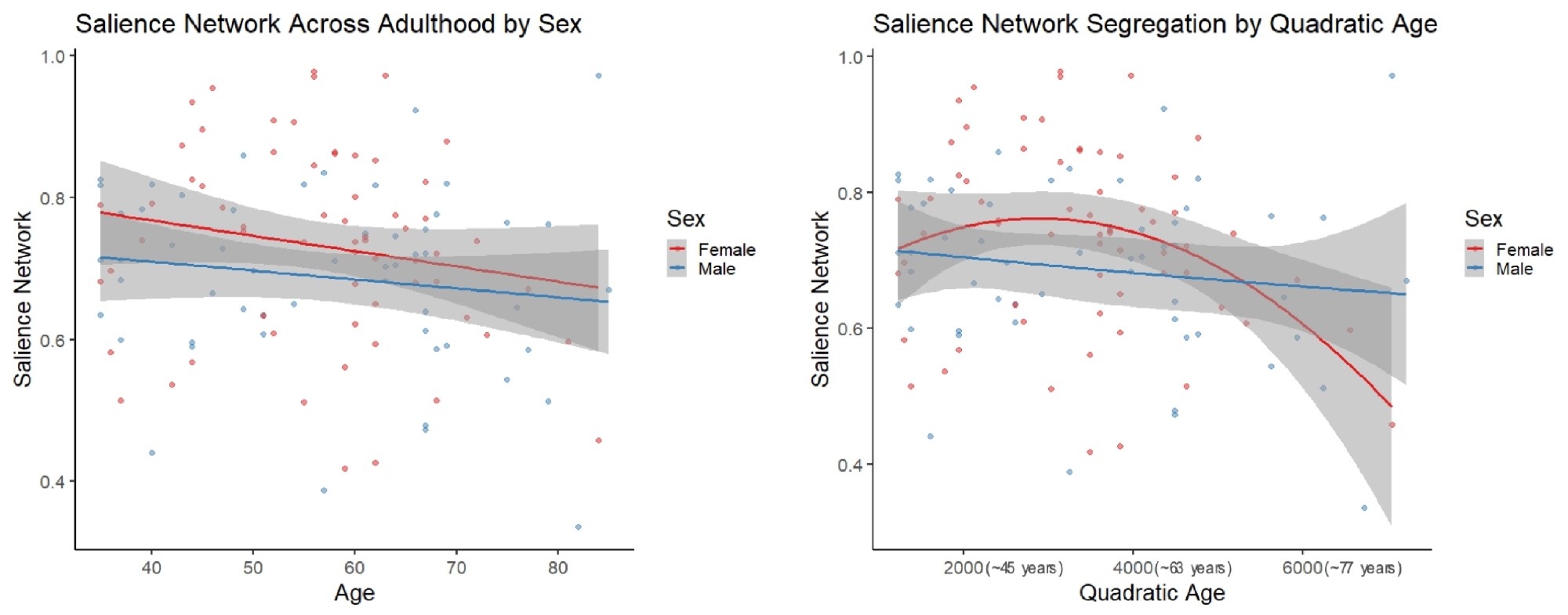
Scatter plots demonstrate the linear (left) and quadratic (right) relationship between Salience network connectivity and age. Parentheses next to quadratic values (right) indicate the approximate square root of the quadratic value for interpretative purposes. The gray superimposed on each colored line depicts the 95% confidence interval in each sex. The quadratic model demonstrated significantly better fit than the linear model for Salience network segregation across adulthood (AIC difference = 11.11).

Comparisons of model fit between linear and quadratic regressions revealed that salience network segregation fits significantly better with a quadratic model as compared to linear in females (AIC difference = 11.11, **Figure 3**). However, all other comparisons of model fit between linear, and quadratic were not statistically significant (see ***Supplementary material* Table 11**).

## 4. Discussion

This study investigated network segregation in aging females in the context of sex steroid hormones. Primarily, we were interested in network segregation in adult females, as sex steroid hormone levels—which undergo dramatic changes in females during mid and later life—may impact brain network properties. Understanding differences in network segregation in females in the context of aging and hormone levels stands to provide a greater understanding around factors contributing to functional differences in aging, particularly given that older females are at greater risk for negative outcomes in later life (42–45). Somewhat surprisingly, we found no significant relationships between sex steroid hormone levels and network segregation levels in adult females. To our knowledge, this is the first study to directly investigate endogenous hormone levels with network segregation and patterns of aging.

In the context of both endogenous hormone levels and exogenous sex hormone treatments, sex hormones have displayed impacts on brain structure and function (e.g., cortical connectivity, subcortical connectivity, and within-network coherence) (11, 46, 47). Given the notable vacillations in hormone levels during distinct reproductive stages (28) and cognitive inefficiencies associated with certain reproductive stages (46–51), we expected to find associations between network segregation and hormone levels. However, females did not exhibit significant associations between progesterone, 17β-estradiol, or testosterone and network segregation. These findings are inconsistent with a recent study of network segregation from our group (23) which demonstrated some differences with reproductive stage, suggesting hormones may play impact network segregation. That is, female reproductive aging is associated with declines in 17β-estradiol and progesterone (10, 42, 52) and Ballard and colleagues (23) demonstrated significant differences between distinct reproductive stages and network segregation (i.e., COTC, DMN, DA, FPTC, and Sa) in females (23). Of note, Ballard and colleagues’ findings may be driven by age as they also found linear relationships between age and network segregation in the aforementioned networks (23). Pritschet and colleagues (2020) demonstrated that estradiol is associated with increasing global efficiency in the DMN and DA networks, whereas progesterone was associated with reduced coherence throughout the brain (46). While Pritschet and colleagues’ findings suggested influences of estradiol and progesterone on network dynamics, it is difficult to directly compare their findings to ours as their study was based on dense sampling in a young female across a menstrual cycle and evaluated a different aspect of network function (46). However, it broadly demonstrates the purported relationship between brain network organization and sex steroid hormones in the female brain. Although we cannot directly compare our findings with this and other studies, network segregation is a proxy for measuring functional organization in the brain and may loosely be interpreted as such.

Lastly, the combined impact (interaction) of hormone levels (i.e., estradiol, progesterone, and testosterone) on network segregation did not reveal a relationship with network segregation in females. We found interactions between estradiol and testosterone in SSH and SSM networks for raw p-values, but these findings did not survive FDR correction. However, this finding agrees with the study by Moffat (2005), in which they note that testosterone can interact not only with androgen receptors, but also with estradiol receptors, and therefore, its administration can, in some cases, parallel the effects of estradiol on the entire nervous system. An important element in understanding the effects of testosterone on the nervous system is that many of its behavioral and anatomical effects occur after it has been converted to its metabolically active derivatives - estradiol or dihydrotestosterone (8).

One consideration beyond the scope of the current study is how stages within a female menstrual cycle or menopausal stage could impact brain connectivity. Syan and colleagues (2017) examined endogenous estradiol, progesterone, and a neuroactive metabolite of progesterone (allopregnanolone) at different menstrual phases and demonstrated an impact of hormone levels in the late luteal phase on resting state connectivity in both cortical and subcortical regions for reproductive aged females (14). As mentioned earlier, Pritschet and colleagues (2020) exhibited an impact of sex hormones on network architecture in tandem with the cycle of a reproductive aged female. Thus, sex steroid hormones have shown a relationship with brain connectivity as it relates to regular menstrual cycle fluctuation. Individual variation exists both in a regular menstrual cycle and menopausal stages (52–54). Notably, hormone levels are not the sole indicator for identifying distinct reproductive stages. Seminal work in categorizing stages in reproductive aging described principal criteria for determining reproductive stage by changes in cycle regularity and days to years since last cycle (52). Endocrine information such as follicle stimulating hormones, antimullerian hormone, and inhibin-B are used to support categorization by changes in cycle (52). We did not examine menstrual cycle, the above specified endocrine information, menopausal stage, or within individual hormone variance in this investigation. As such, the nature of our analyses and scope of our study may not capture the complexity of female network segregation.

In the context of aging, females did display a significant quadratic relationship between age and network segregation for the CBBG and Sa networks. In both cases, segregation was still increasing through adulthood and highest in midlife with a downturn thereafter. Patterns of connectivity between the cerebellum and basal ganglia have been positively linked to cognitive performance and self-reported balance confidence in older adults (26, 55). Further, in their review Diedrichsen and colleagues (2009) (29) implicated the relationship between the cerebellum and basal ganglia as critical for modulating cortical functions such as cognition and relying on subcortical processes. We can see in **Figure 3,** that both females and males demonstrate an inverted “U-shaped” decline in CBBG network segregation across the span of aging adults. Functionally this pattern in females may be related to the drop in cognition and decreased balance that females experience above and beyond males in aging (42, 46), though we should note that this is speculative as we did not include any behavioral analyses here.

The quadratic age relationship with Sa network segregation in females was somewhat consistent with work from Chan and colleagues (2014), where they showed a significant quadratic age relationship across female and male participants in “association systems” which included the Sa network, but was aggregated with DMN, FPTC, VA, COTC, and DA (32). The Sa network has been associated with “conscious integration of autonomic feedback and responses with internal goals and environmental demands” (56). This network has also been conceptualized as an “integral hub” for facilitating communication between the DMN and central executive networks in a triple network model (57). Notably, dysfunction in the Sa network has been linked to reduced cognitive performance in older adults (58) and increased connectivity in this network has been linked to Alzheimer’s disease (59). The quadratic relationship seen in females, but not males (see **Figure 3**), could similarly be highlighting cognitive inefficiencies that have been demonstrated during advanced aging in females (46).

It is important to emphasize that we did not find significant quadratic age relationships in any additional networks for females, nor any networks in males in our exploratory analyses. These findings are consistent with Lee and colleagues’ (2016) study that examined rs-fMRI graph network analysis in the DMN, Sa and Central Executive network and a quadratic age relationship in both groups of “good” and “poor” cognitive performers (60). While Lee and colleagues (2016) assessed components of both network segregation and network integration (i.e., global efficiency, local efficiency, betweenness centrality, connectivity strength, and nodal degree), their analyses notably varied from ours in that they controlled for gender (60). That is, the relationship between quadratic age and segregation in certain networks may be driven by inherent hormonal differences between sexes, but as gender was controlled for in their study that relationship was left unexplored (60). Thus, the methodological differences may explain the differences in outcomes relative to what we report here.

Comparisons of model fit demonstrated the quadratic model as a substantially better fit for Sa network segregation and age in females; however, no additional quadratic age models displayed a significantly better fit in our analyses. To our knowledge, associations between quadratic age and network segregation have not been otherwise evaluated or reported. We would suggest that this may be a useful area of investigation in future work. We did not however find linear relationships between age and network segregation when examining females or males after FDR correction. Similarly, Grady and colleagues (2016), did not see linear relationships with age and network segregation, although their examination was specific to the DMN, DA, and FPTC networks (61). As stated, earlier Chan and colleagues (2014) found a significant quadratic age relationship in association systems (32); however, these systems were also significantly linearly associated with age. The same study also revealed linear age relationships with network segregation in “sensory-motor systems” which aggregated Hand somato-motor, Visual, Mouth somato-motor, and Auditory within-network segregations (62). While we did not replicate Chan and colleagues’ (2014) findings, we also did not aggregate networks for analysis in the same fashion. Further there are also differences in the data used for analyses with respect to both sample size and length of scans. That is, their sample size was twice as large as our study and scan length was about 5 minutes per participant. Of note, our approach to data collection (guided by recent scientific advancement and literature) often produces more reliable data (63) as we collected 24 total minutes of resting state scans, alternating from anterior to posterior slice collection. Thus, Chan and colleagues’ (2014) investigation of network segregation in sensory-motor systems may not be appropriate for drawing direct comparisons to our data given methodological advancements in recent years resulting in differences in data collection parameters. Additional studies evaluating linear and quadratic relationships by sex are warranted, particularly with larger samples that include longer resting state acquisition times such as that used here.

Overall, we found quadratic age relationships for females in CBBG and Sa network segregation. We did not find significant impacts of individual or combined hormone levels on within-network segregation in the networks of interest. Future research would benefit from examining inter-individual differences over time to gain more insight into subtle influences of endogenous hormones. Another relevant area of research would be examining individual differences of network segregation over time in those taking hormonal contraceptives or receiving hormone replacement therapy to better understand the dynamic interplay between sex steroid hormones and brain network organization.

### Limitations

There are several limitations relevant to the present investigation. Namely, the variability and collection of hormone assays and the cross-sectional nature of these data. It is important to recognize that hormone levels can vary almost as much within a naturally cycling female as they can between naturally cycling females (53, 54). Furthermore, the hormone assay was collected on a different day than the scan as freezing the sample at a particular temperature is required immediately, and our study design incorporated a delay to allow for additional measures of activity and questionnaire completion (not related to the research questions presented here). The large variance of hormone levels within a cycle implies that levels could change in a matter of days for reproductive age females. Thus, given the age range of our female sample, the offset between the hormone sample and the scan may impact these relationships for those that still have regular menstrual cycles. While this is only a small subset of this sample, it is an important limitation, nonetheless. Hormone levels vary within and between naturally cycling females so examining sex hormone levels in the same participant over time would account for some individual variance in hormone fluctuations. Lastly, our sex hormone analyses did not account for age. This was intentional as preliminary analyses demonstrated age was significantly correlated with female estradiol and progesterone (*p*<.01) and testosterone maintains a relatively consistent level throughout for mid- to older- age females (64). Thus, incorporating age into hormone-based analyses would account for very similar variance to sex hormone levels.

## Conclusion

This study demonstrated a quadratic relationship between aging and network segregation for the CBBG and Sa networks in females. In both cases, segregation was still increasing through adulthood and highest in midlife with a downturn thereafter. These networks are functionally related to cognitive performance, balance, and integrating autonomic feedback in response to environmental demands (26, 32, 55).

Prior research has shown that the sex hormones can regulate neurogenesis, inflammatory processes, impact network segregation, and may also play a role in regulating cognitive and affective processes, mainly in the aging process (1, 14). Furthermore, the notable variability in hormone levels in certain reproductive stages and cognitive deficits associated with specific reproductive stages did not have an impact on network segregation as was expected (46–51).

Future studies could focus on examining participants longitudinally and pairing these types of data with behavioral outcomes.

## Acknowledgment

This work was supported by R01AG065010 to J.A.B. This work was further supported by the Texas Virtual Data Library (ViDaL), a high-performance cluster, funded by the Texas A&M University Research Development fund. In this cluster the imaging analyses for the current work were carried out using the resources provided by the Texas A&M High Performance Research Computing organization.

