## Supplemental Tables 1-13 for "Network Segregation in Aging Females and Evaluation of the Impact of Sex Steroid Hormones"

### ***Supplementary material***

**Supplementary Table 1.** This table presents results from linear regressions for estradiol and network segregation in males. Raw *p*-values are listed, and FDR correction was only performed if raw *p* -value was <.05.

| Male Estradiol Linear Associations with Network Segregation | | | | | | |
|---|---|---|---|---|---|---|
| | **Estradiol $\beta$ coefficient** | **Raw *p*-value** | **$R^2$** | **Adjusted $R^2$** | **Residual Std. Error (df = 48)** | **F Statistic (df = 1; 48)** |
| **Au** | 0.011 | 0.628 | 0.005 | -0.016 | 0.092 | 0.239 |
| **CBBG** | -0.01 | 0.817 | 0.001 | -0.02 | 0.168 | 0.054 |
| **COTC** | -0.362 | 0.466 | 0.011 | -0.009 | 1.966 | 0.541 |
| **DA** | -0.008 | 0.783 | 0.002 | -0.019 | 0.12 | 0.077 |
| **DM** | -0.01 | 0.772 | 0.002 | -0.019 | 0.136 | 0.085 |
| **FPTC** | -0.059 | 0.102 | 0.055 | 0.035 | 0.142 | 2.789 |
| **Sa** | 0.0003 | 0.994 | 0 | -0.021 | 0.132 | 0.0001 |
| **SSH** | -0.006 | 0.835 | 0.001 | -0.02 | 0.111 | 0.044 |
| **SSM** | 0.01 | 0.571 | 0.007 | -0.014 | 0.067 | 0.327 |
| **Vi** | 0.026 | 0.342 | 0.019 | -0.002 | 0.108 | 0.922 |
| **VA** | 0.009 | 0.832 | 0.001 | -0.02 | 0.165 | 0.046 |

**Supplementary Table 2.** This table presents results from linear regressions for progesterone and network segregation in males. Raw *p*-values are listed, and FDR correction was only performed if raw *p* -value was <.05.

| **Male Progesterone Linear Associations with Network Segregation** | | | | | | |
|---|---|---|---|---|---|---|
| | **Progesterone $\beta$ coefficient** | **Raw *p*-value** | **$R^2$** | **Adjusted $R^2$** | **Residual Std. Error (df = 48)** | **F Statistic (df = 1; 48)** |
| **Au** | 0.0004 | 0.240 | 0.029 | 0.008 | 0.101 | 1.416 |
| **CBBG** | -0.0002 | 0.686 | 0.003 | -0.017 | 0.166 | 0.166 |
| **COTC** | 0.007 | 0.267 | 0.026 | 0.005 | 1.922 | 1.264 |
| **DA** | 0.00002 | 0.951 | 0.0001 | -0.021 | 0.12 | 0.004 |
| **DM** | 0.0002 | 0.723 | 0.003 | -0.018 | 0.141 | 0.128 |
| **FPTC** | -0.0004 | 0.388 | 0.016 | -0.005 | 0.146 | 0.76 |
| **Sa** | 0.0001 | 0.792 | 0.001 | -0.019 | 0.131 | 0.071 |
| **SSH** | -0.0001 | 0.754 | 0.002 | -0.019 | 0.112 | 0.099 |
| **SSM** | 0.0002 | 0.381 | 0.016 | -0.004 | 0.067 | 0.782 |
| **Vi** | 0.0004 | 0.205 | 0.033 | 0.013 | 0.108 | 1.656 |
| **VA** | 0.00003 | 0.955 | 0.0001 | -0.021 | 0.167 | 0.003 |

**Supplementary Table 3.** This table presents results from linear regressions for progesterone and network segregation in males. Raw *p*-values are listed, and FDR correction was only performed if raw *p* -value was <.05.

| Male Testosterone Linear Associations with Network Segregation | | | | | | |
|---|---|---|---|---|---|---|
| | **Testosterone $\beta$ coefficient** | **Raw *p*-value** | **$R^2$** | **Adjusted $R^2$** | **Residual Std. Error (df = 49)** | **F Statistic (df = 1; 49)** |
| **Au** | 0.0002 | 0.493 | 0.01 | -0.011 | 0.1 | 0.478 |
| **CBBG** | -0.0004 | 0.423 | 0.013 | -0.007 | 0.165 | 0.656 |
| **COTC** | 0.001 | 0.867 | 0.001 | -0.02 | 1.947 | 0.029 |
| **DA** | -0.0001 | 0.850 | 0.001 | -0.02 | 0.12 | 0.036 |
| **DM** | -0.00001 | 0.976 | 0.00002 | -0.02 | 0.141 | 0.001 |
| **FPTC** | -0.0003 | 0.473 | 0.011 | -0.01 | 0.145 | 0.525 |
| **Sa** | 0.0001 | 0.816 | 0.001 | -0.019 | 0.13 | 0.055 |
| **SSH** | -0.0003 | 0.354 | 0.018 | -0.002 | 0.11 | 0.879 |
| **SSM** | 0.0001 | 0.781 | 0.002 | -0.019 | 0.066 | 0.078 |
| **Vi** | 0.0004 | 0.253 | 0.027 | 0.007 | 0.106 | 1.342 |
| **VA** | 0.0003 | 0.517 | 0.009 | -0.012 | 0.165 | 0.428 |

**Supplementary Table 4.** This table presents results from linear regressions for estradiol and network segregation across male and female participants. Raw *p*-values are listed, and FDR correction was only performed if raw *p* -value was <.05.

| Estradiol Linear Associations with Network Segregation | | | | | | |
|---|---|---|---|---|---|---|
| | **Estradiol $\beta$ coefficient** | **Raw *p*-value** | **$R^2$** | **Adjusted $R^2$** | **Residual Std. Error (df = 105)** | **F Statistic (df = 1; 105)** |
| **Au** | 0.006 | 0.717 | 0.001 | -0.008 | 0.098 | 0.132 |
| **CBBG** | 0.017 | 0.538 | 0.004 | -0.006 | 0.161 | 0.384 |
| **COTC** | 0.45 | 0.446 | 0.006 | -0.004 | 3.42 | 0.586 |
| **DA** | -0.012 | 0.557 | 0.003 | -0.006 | 0.114 | 0.348 |
| **DM** | 0.01 | 0.688 | 0.002 | -0.008 | 0.142 | 0.163 |
| **FPTC** | -0.017 | 0.472 | 0.005 | -0.005 | 0.137 | 0.522 |
| **Sa** | 0.028 | 0.239 | 0.013 | 0.004 | 0.137 | 1.402 |
| **SSH** | -0.006 | 0.743 | 0.001 | -0.008 | 0.11 | 0.109 |
| **SSM** | -0.007 | 0.601 | 0.003 | -0.007 | 0.073 | 0.276 |
| **Vi** | 0.009 | 0.683 | 0.002 | -0.008 | 0.124 | 0.168 |
| **VA** | -0.011 | 0.776 | 0.001 | -0.009 | 0.219 | 0.081 |

**Supplementary Table 5.** This table presents results from linear regressions for estradiol and network segregation across male and female participants. Raw *p*-values and FDR corrected *p*-values are included. Asterisks indicate significance at *p*<.05* for FDR corrected values. Only FDR corrected values are interpreted as significant.

| Progesterone Linear Associations with Network Segregation | | | | | | | |
|---|---|---|---|---|---|---|---|
| | **Progesterone β coefficient** | **Raw *p*-value** | **FDR Corrected *p*-value** | **$R^2$** | **Adjusted $R^2$** | **Residual Std. Error (df = 109)** | **F Statistic (df = 1; 109)** |
| **Au** | 0.0002 | 0.187 | 0.411 | 0.016 | 0.007 | 0.105 | 1.767 |
| **CBBG** | 0.0002 | 0.245 | 0.449 | 0.012 | 0.003 | 0.165 | 1.37 |
| **COTC** | 0.014 | 0.001 | 0.011* | 0.101 | 0.093 | 3.142 | 12.257 |
| **DA** | 0.0001 | 0.563 | 0.746 | 0.003 | -0.006 | 0.113 | 0.337 |
| **DM** | 0.0003 | 0.163 | 0.411 | 0.018 | 0.009 | 0.142 | 1.98 |
| **FPTC** | 0.0002 | 0.163 | 0.411 | 0.018 | 0.009 | 0.135 | 1.973 |
| **Sa** | 0.0003 | 0.056 | 0.308 | 0.033 | 0.024 | 0.133 | 3.756 |
| **SSH** | 0.00004 | 0.799 | 0.836 | 0.001 | -0.009 | 0.113 | 0.066 |
| **SSM** | 0.00005 | 0.610 | 0.746 | 0.002 | -0.007 | 0.075 | 0.263 |
| **Vi** | 0.0001 | 0.435 | 0.684 | 0.006 | -0.004 | 0.122 | 0.615 |
| **VA** | -0.0001 | 0.836 | 0.836 | 0.0004 | -0.009 | 0.212 | 0.043 |

**Supplementary Table 6.** This table presents results from linear regressions for estradiol and network segregation across male and female participants. Raw *p*-values are listed, and FDR correction was only performed if raw *p* -value was <.05.

| **Testosterone Linear Associations with Network Segregation** | | | | | | |
|---|---|---|---|---|---|---|
| | **Testosterone $\beta$ coefficient** | **Raw *p*-value** | **$R^2$** | **Adjusted $R^2$** | **Residual Std. Error (df = 111)** | **F Statistic (df = 1; 111)** |
| **Au** | -0.0002 | 0.288 | 0.01 | 0.001 | 0.105 | 1.144 |
| **CBBG** | -0.0002 | 0.433 | 0.006 | -0.003 | 0.165 | 0.62 |
| **COTC** | -0.004 | 0.519 | 0.004 | -0.005 | 3.331 | 0.419 |
| **DA** | -0.0002 | 0.324 | 0.009 | -0.0001 | 0.112 | 0.983 |
| **DM** | -0.0001 | 0.822 | 0.0005 | -0.009 | 0.143 | 0.051 |
| **FPTC** | -0.0002 | 0.305 | 0.009 | 0.001 | 0.134 | 1.062 |
| **Sa** | -0.0002 | 0.440 | 0.005 | -0.004 | 0.135 | 0.601 |
| **SSH** | -0.0002 | 0.358 | 0.008 | -0.001 | 0.112 | 0.855 |
| **SSM** | -0.0001 | 0.498 | 0.004 | -0.005 | 0.074 | 0.464 |
| **Vi** | -4E-05 | 0.840 | 0.0004 | -0.009 | 0.121 | 0.041 |
| **VA** | 0 | 0.998 | 0 | -0.009 | 0.212 | 0.00001 |

**Supplementary Table 7.** This table presents results from linear regressions for linear age and network segregation in males. Raw *p*-values are listed, and FDR correction was only performed if raw *p* -value was <.05.

| Male Linear Age Associations with Network Segregation | | | | | | |
|---|---|---|---|---|---|---|
| | **Age $\beta$ coefficient** | **Raw *p*-value** | **$R^2$** | **Adjusted $R^2$** | **Residual Std. Error (df = 53)** | **F Statistic (df = 1; 53)** |
| **Au** | -0.001 | 0.389 | 0.014 | -0.005 | 0.098 | 0.755 |
| **CBBG** | -0.001 | 0.538 | 0.007 | -0.012 | 0.164 | 0.385 |
| **COTC** | -0.022 | 0.207 | 0.03 | 0.012 | 1.882 | 1.635 |
| **DA** | -0.001 | 0.533 | 0.007 | -0.011 | 0.117 | 0.395 |
| **DM** | 0.001 | 0.606 | 0.005 | -0.014 | 0.14 | 0.269 |
| **FPTC** | -0.001 | 0.644 | 0.004 | -0.015 | 0.143 | 0.217 |
| **Sa** | -0.001 | 0.286 | 0.022 | 0.003 | 0.127 | 1.166 |
| **SSH** | -0.0005 | 0.638 | 0.004 | -0.015 | 0.109 | 0.224 |
| **SSM** | -0.0005 | 0.452 | 0.011 | -0.008 | 0.068 | 0.575 |
| **Vi** | -0.002 | 0.081 | 0.057 | 0.039 | 0.114 | 3.184 |
| **VA** | -0.0004 | 0.796 | 0.001 | -0.018 | 0.166 | 0.068 |

**Supplementary Table 8.** This table presents results from linear regressions for age and network segregation across male and female participants. Raw *p*-values and FDR corrected *p*-values are included. Asterisks indicate significance at *p*<.05* for FDR corrected values. Only FDR corrected values are interpreted as significant.

| Linear Age Associations with Network Segregation | | | | | | | |
|---|---|---|---|---|---|---|---|
| | **Age $\beta$ coefficient** | **Raw *p*-value** | **FDR Corrected *p*-value** | **$R^2$** | **Adjusted $R^2$** | **Residual Std. Error (df = 117)** | **F Statistic (df = 1; 117)** |
| **Au** | -0.001 | 0.074 | 0.178 | 0.027 | 0.019 | 0.103 | 3.273 |
| **CBBG** | -0.002 | 0.106 | 0.194 | 0.022 | 0.014 | 0.165 | 2.655 |
| **COTC** | -0.047 | 0.040 | 0.178 | 0.036 | 0.028 | 3.214 | 4.337 |
| **DA** | -0.001 | 0.325 | 0.397 | 0.008 | -0.0002 | 0.111 | 0.977 |
| **DM** | -0.0002 | 0.879 | 0.879 | 0.0002 | -0.008 | 0.143 | 0.023 |
| **FPTC** | -0.001 | 0.212 | 0.333 | 0.013 | 0.005 | 0.134 | 1.581 |
| **Sa** | -0.002 | 0.079 | 0.178 | 0.026 | 0.018 | 0.133 | 3.154 |
| **SSH** | -0.001 | 0.383 | 0.421 | 0.007 | -0.002 | 0.111 | 0.77 |
| **SSM** | -0.001 | 0.073 | 0.178 | 0.027 | 0.019 | 0.073 | 3.282 |
| **Vi** | -0.002 | 0.081 | 0.178 | 0.026 | 0.018 | 0.123 | 3.115 |
| **VA** | -0.002 | 0.283 | 0.389 | 0.01 | 0.001 | 0.211 | 1.168 |

**Supplementary Table 9.** This table presents results from quadratic regressions for age and network segregation in male participants. Raw *p*-values and FDR corrected *p*-values are included. Asterisks indicate significance at *p*<.05* for FDR corrected values. Only FDR corrected values are interpreted as significant.

| Male Quadratic Age Associations with Network Segregation | | | | | | | |
|---|---|---|---|---|---|---|---|
| | **Quadratic Age β coefficient** | **Raw *p*-value** | **FDR Corrected *p*-value** | **$R^2$** | **Adjusted $R^2$** | **Residual Std. Error (df = 52)** | **F Statistic (df = 2; 52)** |
| **Au** | -0.0001 | 0.084 | 0.462 | 0.07 | 0.034 | 0.096 | 1.954 |
| **CBBG** | -0.0003 | 0.020 | 0.220 | 0.107 | 0.073 | 0.157 | 3.116 |
| **COTC** | -0.0002 | 0.904 | 0.962 | 0.03 | -0.007 | 1.899 | 0.81 |
| **DA** | 0.00001 | 0.941 | 0.962 | 0.008 | -0.031 | 0.119 | 0.197 |
| **DM** | -0.0001 | 0.409 | 0.962 | 0.018 | -0.02 | 0.141 | 0.481 |
| **FPTC** | 0.0001 | 0.495 | 0.962 | 0.013 | -0.025 | 0.143 | 0.344 |
| **Sa** | -0.00001 | 0.922 | 0.962 | 0.022 | -0.016 | 0.128 | 0.577 |
| **SSH** | -0.00003 | 0.731 | 0.962 | 0.007 | -0.032 | 0.11 | 0.17 |
| **SSM** | 0.00001 | 0.756 | 0.962 | 0.013 | -0.025 | 0.068 | 0.332 |
| **Vi** | 0 | 0.962 | 0.962 | 0.057 | 0.02 | 0.115 | 1.563 |
| **VA** | -0.00004 | 0.713 | 0.962 | 0.004 | -0.034 | 0.167 | 0.102 |

**Supplementary Table 10.** This table presents results from quadratic regressions for age and network segregation across male and female participants. Raw *p*-values and FDR corrected *p*-values are included. Asterisks indicate significance at *p*<.05* for FDR corrected values. Only FDR corrected values are interpreted as significant.

| Quadratic Age Associations with Network Segregation | | | | | | | |
|---|---|---|---|---|---|---|---|
| | **Quadratic Age $\beta$ coefficient** | **Raw *p*-value** | **FDR Corrected *p*-value** | **$R^2$** | **Adjusted $R^2$** | **Residual Std. Error (df = 116)** | **F Statistic (df = 2; 116)** |
| **Au** | -0.0002 | 0.003 | 0.017* | 0.101 | 0.085 | 0.099 | 6.515 |
| **CBBG** | -0.0003 | 0.001 | 0.011* | 0.117 | 0.101 | 0.157 | 7.660 |
| **COTC** | -0.002 | 0.288 | 0.390 | 0.045 | 0.029 | 3.212 | 2.741 |
| **DA** | -0.0001 | 0.236 | 0.390 | 0.02 | 0.003 | 0.111 | 1.203 |
| **DM** | -0.0001 | 0.072 | 0.198 | 0.028 | 0.011 | 0.142 | 1.664 |
| **FPTC** | -0.00003 | 0.637 | 0.637 | 0.015 | -0.002 | 0.134 | 0.897 |
| **Sa** | -0.0001 | 0.034 | 0.125 | 0.063 | 0.047 | 0.131 | 3.932 |
| **SSH** | -0.0001 | 0.096 | 0.211 | 0.03 | 0.013 | 0.11 | 1.805 |
| **SSM** | -0.00004 | 0.317 | 0.390 | 0.036 | 0.019 | 0.073 | 2.148 |
| **Vi** | -0.0001 | 0.319 | 0.390 | 0.034 | 0.018 | 0.123 | 2.061 |
| **VA** | -0.0001 | 0.472 | 0.519 | 0.014 | -0.003 | 0.212 | 0.842 |

**Supplementary Table 11.** Superior model fit would be determined by a difference of |10| between linear and quadratic models. Asterisk indicates superior fit in a quadratic model as compared to a linear model for age and network segregation.

| **Female Linear and Quadratic Age Relationships with Network Segregation AIC Model Fit Comparisons** | | | |
|---|---|---|---|
| | **Female Linear AIC** | **Female Quadratic AIC** | **Female Difference (Linear AIC-Quad AIC)** |
| **Au** | -106.723 | -111.224 | 4.502 |
| **CBBG** | -44.239 | -53.560 | 9.321 |
| **COTC** | 371.517 | 372.737 | -1.220 |
| **DA** | -105.371 | -106.819 | 1.448 |
| **DM** | -63.105 | -65.445 | 2.339 |
| **FPTC** | -81.679 | -83.478 | 1.800 |
| **Sa** | -67.344 | -78.458 | 11.114* |
| **SSH** | -96.195 | -98.919 | 2.724 |
| **SSM** | -146.535 | -146.629 | 0.094 |
| **Vi** | -77.789 | -77.762 | -0.028 |
| **VA** | 3.374 | 5.046 | -1.672 |

**Supplementary Table 12.** This table exhibits results from linear regressions for hormone interactions and network segregation in males. Raw *p*-values and FDR corrected *p*-values are included. FDR correction was only performed if raw *p* -value was <.05. There were no significant findings after FDR correction.

| Male Hormone Level Interactions with Network Segregation | | | | | | | | | | | |
|---|---|---|---|---|---|---|---|---|---|---|---|
| | **Estradiol by Progesterone** (E*P) ***β* coefficient** | **Raw E*P *p*-value** | **Estradiol by Testosterone** (E*T) ***β* coefficient** | **Raw E*T *p*-value** | **Progesterone by Testosterone** (P*T) ***β* coefficient** | **Raw P*T *p*-value** | **FDR Corrected *p*-value for Estradiol by Progesterone** | **$R^2$** | **Adjusted $R^2$** | **Residual Std. Error (df = 39)** | **F Statistic (df = 6; 39)** |
| **Au** | -0.0004 | 0.592 | 0.001 | 0.299 | 0 | 0.634 | 0.627 | 0.07 | -0.074 | 0.097 | 0.486 |
| **CBBG** | -0.001 | 0.627 | 0.002 | 0.463 | 0 | 0.955 | 0.627 | 0.04 | -0.108 | 0.176 | 0.272 |
| **COTC** | -0.033 | 0.041 | 0.016 | 0.468 | -0.0003 | 0.123 | 0.226 | 0.278 | 0.167 | 1.818 | 2.500 |
| **DA** | -0.001 | 0.361 | 0.002 | 0.242 | -0.00002 | 0.153 | 0.627 | 0.072 | -0.071 | 0.127 | 0.501 |
| **DM** | -0.001 | 0.549 | 0.001 | 0.496 | -0.00001 | 0.603 | 0.627 | 0.037 | -0.111 | 0.146 | 0.25 |
| **FPTC** | -0.003 | 0.032 | 0.003 | 0.113 | -0.00001 | 0.410 | 0.226 | 0.168 | 0.04 | 0.147 | 1.309 |
| **Sa** | 0.001 | 0.622 | -0.0002 | 0.924 | -0.00001 | 0.537 | 0.627 | 0.024 | -0.126 | 0.142 | 0.163 |
| **SSH** | -0.001 | 0.340 | 0.002 | 0.252 | 0 | 0.752 | 0.627 | 0.101 | -0.037 | 0.116 | 0.733 |
| **SSM** | -0.001 | 0.187 | 0.001 | 0.469 | -0.00001 | 0.251 | 0.611 | 0.12 | -0.016 | 0.067 | 0.884 |
| **Vi** | -0.001 | 0.222 | 0.001 | 0.328 | 0 | 0.694 | 0.611 | 0.074 | -0.068 | 0.114 | 0.523 |
| **VA** | -0.001 | 0.467 | 0.0004 | 0.834 | -0.00001 | 0.471 | 0.627 | 0.054 | -0.092 | 0.176 | 0.368 |

**Supplementary Table 13.** This table exhibits results from linear regressions for hormone interactions and network segregation in across females and males. Raw *p*-values are included. FDR correction was only performed if raw *p* -value was <.05.

| Hormone Level Interactions with Network Segregation | | | | | | | | | | |
|---|---|---|---|---|---|---|---|---|---|---|
| | **Estradiol by Progesterone** (E*P) ***β* coefficient** | **Raw E*P *p*-value** | **Estradiol by Testosterone** (E*T) ***β* coefficient** | **Raw E*T *p*-value** | **Progesterone by Testosterone** (P*T) ***β* coefficient** | **Raw P*T *p*-value** | **$R^2$** | **Adjusted $R^2$** | **Residual Std. Error (df = 94)** | **F Statistic (df = 6; 94)** |
| **Au** | -0.0001 | 0.681 | -0.0001 | 0.917 | 0 | 0.484 | 0.059 | -0.002 | 0.1 | 0.974 |
| **CBBG** | 0.001 | 0.199 | -0.0001 | 0.929 | 0 | 0.704 | 0.045 | -0.016 | 0.163 | 0.744 |
| **COTC** | -0.016 | 0.148 | -0.015 | 0.410 | 0.0001 | 0.618 | 0.165 | 0.111 | 3.241 | 3.091 |
| **DA** | -0.0005 | 0.219 | 0.0004 | 0.564 | 0 | 0.590 | 0.041 | -0.021 | 0.116 | 0.662 |
| **DM** | -0.0002 | 0.623 | 0 | 0.999 | 0 | 0.959 | 0.029 | -0.033 | 0.145 | 0.461 |
| **FPTC** | -0.001 | 0.267 | -0.0005 | 0.510 | 0 | 0.872 | 0.091 | 0.033 | 0.136 | 1.561 |
| **Sa** | 0.0003 | 0.490 | -0.0004 | 0.578 | 0 | 0.938 | 0.056 | -0.004 | 0.138 | 0.936 |
| **SSH** | 0.00002 | 0.964 | -0.0002 | 0.781 | 0 | 0.723 | 0.022 | -0.04 | 0.114 | 0.357 |
| **SSM** | -0.0003 | 0.253 | 0.0001 | 0.735 | 0 | 0.765 | 0.03 | -0.032 | 0.074 | 0.486 |
| **Vi** | -0.001* | 0.084 | 0.0005 | 0.483 | 0 | 0.917 | 0.043 | -0.018 | 0.127 | 0.705 |
| **VA** | 0.0001 | 0.881 | -0.0004 | 0.742 | 0 | 0.940 | 0.004 | -0.06 | 0.226 | 0.057 |
